# Structural and Cognitive Solutions to Prevent Group Fragmentation in Group-Living Species

**DOI:** 10.1101/2022.12.13.520310

**Authors:** R.I.M. Dunbar

**Author notes:** [ ].

## Abstract

Group-living is one of the six major evolutionary transitions. However, group-living creates stresses that naturally cause group fragmentation, and hence loss of the benefits that group-living provides. How species that live in large groups counteract these forces is not well understood. I analyse comparative data on grooming networks from a large sample of primate species and show that two different social grades can be differentiated in terms of network size and structure. I show that living in large, stable groups involves a combination of increased investment in bonding behaviours (made possible by a dietary adjustment) and the evolution of neuronally expensive cognitive skills of the kind known to underpin social relationships in humans. The first allows the stresses created by these relationships to be defused; the second allows large numbers of weak relationships to be managed, creating a form of multilevel sociality based on strong versus weak ties similar to that found in human social networks.

---

Group-living is one of the six major evolutionary transitions [1]. However, living in groups creates ecological and psychological stresses that, all else equal, impose a limit on group size [2]. In primates, the most intensely social of all mammals, these stresses are so severe that they would limit group size to ~15 individuals [2]. Yet some primate species live in stable groups that number 40-100 individuals in size [2–3]. In order to live in such large groups, animals need to find ways to mitigate these stresses. Herd-forming species (most artiodactyls, mysticete cetaceans) avoid this problem by allowing individuals to join and leave on the basis of the momentary costs and benefits of being in the group, as specified by classic joiner-leaver models [4]. In these species, individuals are effectively anonymous and personalised relationships are rare. In contrast, species that live in stable, bonded social groups (anthropoid primates, equids, tylopods, some suricates, elephants, delphinids) [1,5–7] exploit personalised (in many cases, life-long) relationships that buffer individuals against the stresses of group-living, thereby allowing them to live together in larger, more stable groups. How they manage this is not known.

In herd-forming species, groups are unstable principally because, at a proximate level, individual activity schedules get out of synchrony: groups fragment and disperse because some individuals stop to rest while others continue foraging [8–12]. The stability of bonded social groups, in contrast, arises from individuals’ concerns not to become separated from their principal social partners [13]. This means they constantly monitor social partners [14] and continually reset their activity priorities so as to ensure that everyone’s schedules remain synchronised. Even so, bonded social groups can become dispersed and fragment when day journeys are long and/or group size is large (*Pan* [15]; *Papio* [12,16–17]; *Theropithecus* [18]).

Not only are anthropoid primates among the most intensely social of all mammals, they also have the largest stable groups, as well as some of the largest herd-like groupings [2]. This makes primates a particularly useful taxon for understanding how animals manage the stresses of living in large groups. In some cases, group coordination is ensured by behavioural mechanisms explicitly designed to maintain group cohesion (e.g. ‘notifying’ rituals whereby a group agrees on the foraging route they will take during the day [19–21]). In addition, social grooming is thought to play a key role, not least because some of the most social species devote as much as 20% of their day to it [22–23]). In both monkeys [24] and humans [25–26], the frequency of social interaction directly determines willingness to provide coalitionary support or other forms of altruistic aid, and creates the focus for the social monitoring that ensures that social partners stay together.

Grooming influences bond formation through its effect on the central endorphin system [27–32]. The hand actions used in grooming, and ‘soft touch’ more generally, activate the highly specialised afferent c-tactile (CT) peripheral nerve system [33–35] that triggers the release of β-endorphins in the brain [32]. In humans, endorphin uptake has been shown to be explicitly involved in social bonding [36–39]. This neural system works in tandem with a second cognitive process based on a separate neural pathway to create a dual process mechanism underpinning social bonding [40]. This second cognitive mechanism is not simply a matter of memory capacity and associative learning, but involves sophisticated forms of meta-cognition such as the capacity to infer abstract rules, analogical reasoning, mentalising and self-control that allow relationships to be managed. These cognitive capacities are unique to the anthropoid primates [41–42] and depend on brain regions (notably Brodman areas BA9/10 in the frontal pole) that are only found in this suborder [42,43].

If each bonded relationship requires, as seems to be the case [24–25,44], the investment of a minimum amount of time to ensure functionality and the time available for social grooming is ecologically limited [45], animals will be forced to choose between (a) dividing their limited available social time equally between all group members (at the cost of having increasingly poorly bonded relationships as group size increases) and (b) focussing their social effort on just a few individuals with whom they can maintain optimally bonded relationships (at the cost of sacrificing relationship quality with all other group members) [7]. Both strategies will result in unstable groups – the one because all bonds will be too weak to prevent individuals wandering away from the group during foraging [8–11], the other because the group will fragment into a set of disconnected subnetworks that are prone to becoming separated. Evidence from primates indicates that groups whose habitats limit the time they have available for social grooming are more likely to fragment during foraging than similar sized groups of the same species that can afford to devote more time to grooming [16].

There are three possible ways primates might solve this dilemma in order to live in larger groups. These focus on structural, behavioural and cognitive solutions. The latter two focus, respectively, on the grooming and cognitive components of the dual process mechanism that underpins social bonding.

The first of these solutions is structural (hypothesis H1). Primate grooming networks naturally partition into two layers: grooming cliques (in network terminology, their degree, defined by the number of grooming partners an individual has) and grooming chains (*n*-cliques, defined as the set of individuals linked together by a chain of such ties, even if they do not individually groom each other) [46–48] (Fig. 1). When individuals cannot afford the time to groom every member of their group to the requisite level, these extended chains of virtual relationships might provide the basis for maintaining spatial cohesion during travel when groups get large [13]. Each individual simply has to monitor its principal grooming partners, and a ‘friends-of-a-friend’ effect will keep the entire group together as an extended chain. In effect, grooming patterns form zones of declining gravitational attraction that radiate outwards around each individual to include some or all of the group, creating a form of gravitational drag (Fig. 1, inset) that holds the set of animals together when their individual personal networks are mapped on top of each other. By reducing the number of fracture lines within the network across which no grooming (or social monitoring) takes place, the likelihood that individuals, or even subsets of individuals, will drift off on their own during foraging is reduced. H1 predicts that the size of these extended grooming chains will increase proportionately to group size, even if the number of individuals groomed remains constant; more importantly, grooming chains should encompass most (if not all) of the adults in a group so as to hold them in a tight structural web. If they do not, it implies that other mechanisms must play a role in bridging the gaps between the subnetworks.

**Fig. 1.**
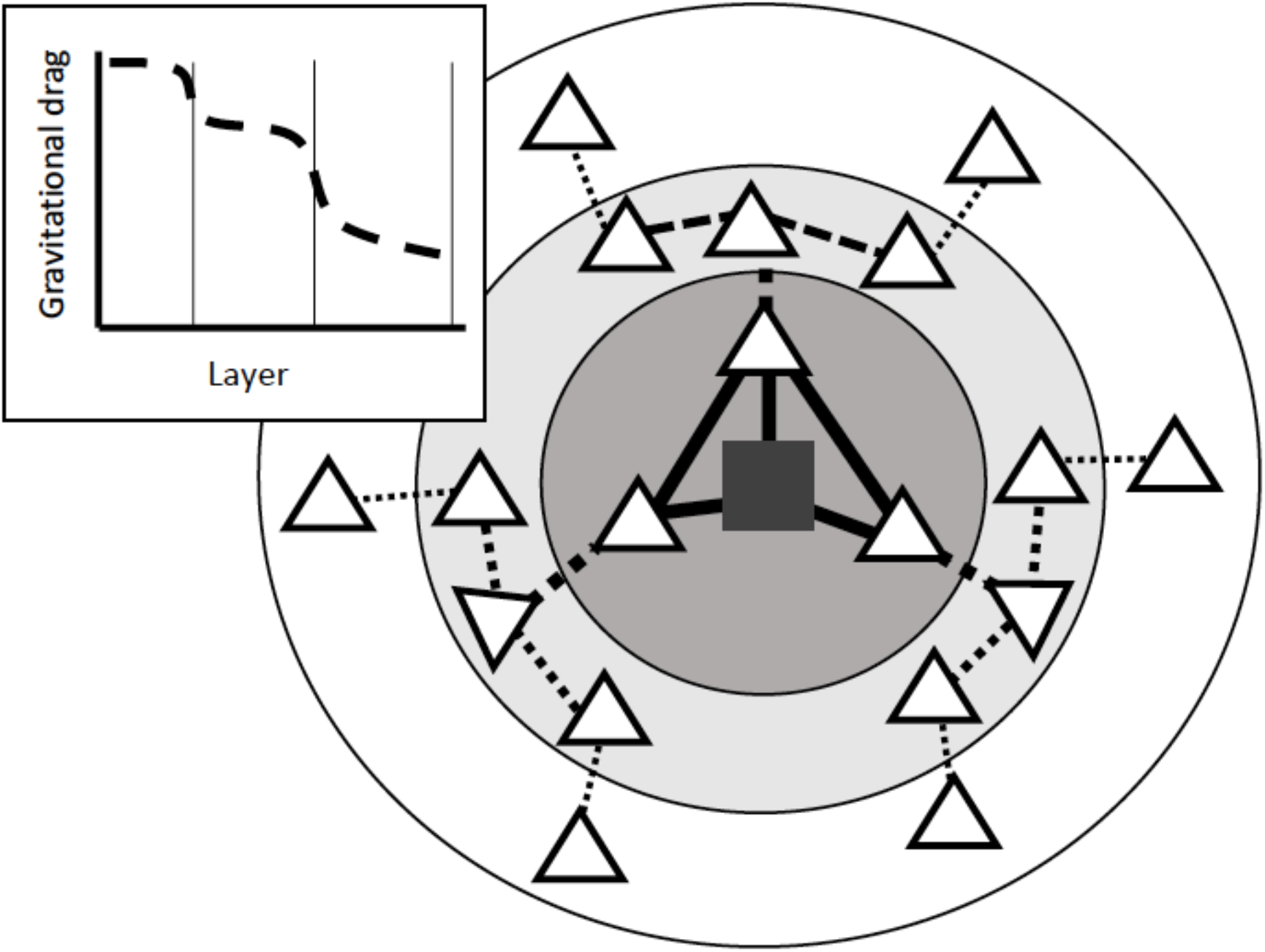
The layered structure of primate and human egocentric social networks, as reflected in grooming patterns. Dark grey square: Ego; white triangles: individual alters. Group members not connected by grooming directly or indirectly to Ego are not shown. Lines connecting triangles indicate grooming relationships, with strength of relationship indicated by width of tie; dashed ties indicate indirect connections (‘friends-of-friends’). The layers of the network are indicated by the shaded circles. Inset: bond strength indicating gravitational drag on dyads willingness to disperse as a function of network layer (i.e. relationship strength).

The second possibility (hypothesis H2) is behavioural. The most pressing issue for group-living mammals is the need to mitigate the stresses incurred from living in close physical proximity with many other individuals. This is best done by devoting disproportionately more grooming time to core allies so as to ensure that these will always be nearby and willing to support each other in any conflicts that arise [24,49–50]. In this way, the benefits of living in a group are retained but the costs are minimised, thereby tilting the cost/benefit ratio in favour of remaining in the group. It may not prevent fragmentation happening altogether, since other factors such as activity desynchrony may still result in groups dispersing when they become very large or travel very long distances [10,12,17]. Nonetheless, it may be enough to make the problem manageable, thereby deferring the point at which groups naturally fragment [2]. H2 thus focusses on the ‘strong ties’ at the centre of each individual’s social network. It predicts that animals will increase the amount of grooming directed to their core grooming partners in proportion to increasing group size in order to reinforce their key alliances, but that the size of both grooming cliques (degree) and grooming chains size will be unrelated to group size (or even reduce in size).

The third possibility is cognitive (hypothesis H3). Animals may be able to mitigate the stresses of group-living by using more sophisticated cognitive mechanisms that allow them to predict and manage others’ behaviour, especially those with whom their do not normally groom. This includes finetuning when and how to respond to others’ threats or spatial incursions, or knowing whom to avoid conflict with because they have more and/or higher rank allies (third party knowledge). This mechanism focusses not on the behavioural mechanisms involved in dyadic social interaction (the focus of hypothesis H2) but on the cognitive mechanisms that allow individuals to manage “weak ties” in the periphery of the network so as to defuse conflict and minimise the risk causing the group to fragment following escalated conflicts. These kinds of high order cognitive skills are associated with executive function and include causal reasoning, analogical reasoning, one trial learning, self-control (inhibition, the ability to defer reward) and mentalising (the capacity to understand others’ intentions), all of which are correlated (both within and between species) with the volume of the brain’s default mode neural network (and hence with brain size) [42–43,51–56]. H3 predicts that species in larger groups will score higher on these cognitive indices than those that live in smaller groups, but not necessarily groom with more individuals or devote more time to grooming.

These are not mutually exclusive alternatives. Rather, they are additive in the sense that they reflect ways of managing the stresses of group-living that differ in their level of cognitive demand. As such, they might represent progressively more sophisticated solutions to successive glass ceilings so as to allow progressively larger groups to be formed in a stepwise manner.

## Methods

Because we are interested in the processes that underpin ties and create networks and, in primates, this is primarily mediated by social grooming, I use only social network data based on allogrooming. Although many published studies of primate social networks use proximity data (either alone or combined with grooming frequencies, sometimes in the form of Silk’s [57] composite social index), proximity is a *consequence* not the cause of ties (see *SI Supplementary Methods*). More importantly, networks based on spatial proximity do not always correlate especially well with networks based on grooming [13,57]. For these reasons, I consider only grooming networks I found data on the distribution of dyadic grooming frequencies for 92 social groups (953 adults) from 36 species (mean = 2.4, range 1-13, groups/species) belonging to 20 genera (mean = 1.8, range 1-7, species/genus) (for details, see *SI Supplementary Methods*). Pair-living species were excluded, since grooming networks are meaningless when there are only two adults. Similarly, grooming with and between immature (pre-puberty) animals were not included.

The grooming matrix for each group was used to calculate two structural indices: the number of adult grooming partners for each individual (its degree) and the number of (adult) individuals connected to each other directly through a grooming relationship or indirectly through a chain of such relationships (an *n*-clique, indexed as all the individuals linked together by an unbroken chain of ties). Degree is a property of the individual; *n*-cliques are subnetworks, and hence a property of the network. Each of these indices was averaged first for each group, then for each species, and finally for each genus.

The mean social group size and number of adult females per group characteristic of a species are taken from [3]. (Issues relating to the definition of social groups are discussed in *SI Supplementary Methods*.) I use four different indices of cognitive ability. Neocortex ratio (the best predictor of social group size as well as of cognitive skills) is taken from [41]; executive function cognition is indexed as the mean of the arcsin transform of the proportion of correct trials on eight standard executive function tasks that test reasoning skills (detour tasks, reversal learning, oddity problems, learning set and delayed reward tasks), from [58]; data on self-control (behavioural inhibition) are from [17] based on a standard A-not-B task; and, finally, data on mentalising competences based on the ‘hide and seek’ game with a human are from [60]. Although data on mentalising competences are available only for three of the genera in the present sample, they are of particular interest because of their implications for animals’ abilities to make correct predictions about others’ intentions. Data on percent of the day devoted to social grooming are taken from [22–23]; *Lemur* and *Eulemur* were excluded because the data given by these sources include autogrooming as well as social grooming. Diet data (percentage of the diet made up of leaves) are from [60].

Data for both individual groups and genus averages are given in online *SI Dataset-1* and *Dataset-2*.

I use *k*-means cluster analysis to determine whether the distribution of species mean group sizes consists of a single homogenous set or two or more distinct grades. Methods for identifying the optimal number of clusters are given in Figs. S1-S5. In most cases, cluster analysis assigns the species of a genus to the same grade; where they do not, I assign all the species to the grade to which the majority of its species are assigned (cf. [41]).

I do not use phylogenetic methods because these cannot identify grades when present in data [2,41]. As it happens, in primates, behavioural and ecological variables have very low or negligible phylogenetic signals [61]; no analysis of primate behaviour and ecology with and without using phylogenetic methods has ever changed any results [62]. Nonetheless, to be sure that significance levels are not inflated by failure to control for phylogenetic effects, all analyses were run separately at group, species and genus levels. If there is no phylogenetic autocorrelation, the results should not differ. In addition, where appropriate, I re-ran some main analyses with phylogenetic correction, but in no case did it change any results (see Fig. S6).

For graphical clarity, all analyses are shown at genus level; the results for species- and group-level analyses are given in the *SI* (see Figs. S7-S10).

## Results

A *k*-means cluster analysis of group size finds a significant partition into two distinct subgroups: genera with small social groups (<25 individuals of all ages) and those with large (>25) groups (genus level: F_1,18_=73.3, p<<0.0001; species level: F_1,34_=84.6, p<<0.0001; group level: F_1,77_=351.8, p<<0.0001) (see *SI*, Figs. S2-S5). Mean group sizes at genus level are 14.9 (range 8.1-24.8, with 5.6±1.96 adult females; N=13 genera; species level mean = 15.7; individual group mean = 14.2) and 36.0 (range 29.3-44.4, with 13.4±1.25 females; N=7; species mean = 40.5; individual group mean = 40.5), respectively. Although all the strepsirrhines and New World monkeys are included in the lower grade and the only ape (*Pan*) in the upper one, Old World monkeys appear in both grades (cutting across taxonomic families). This suggests a functional division rather than a strictly phyletic one. In all three analyses, *Semnopithecus*, *Piliocolobus*, *Chlorocebus*, *Erythrocebus*, *Macaca*, *Theropithecus*, *Papio* and *Pan* are assigned to the upper grade, and *Lemur*, *Eulemur*, *Callithrix*, *Cebus*, *Sapajus, Saimiri, Alouatta*, *Ateles*, *Trachypithecus*, *Nasalis*, *Colobus* and *Cercopithecus* to the lower grade.

### Network structure (hypothesis H1)

Fig. 2 plots mean *n*-clique size against mean degree for the different genera as a function of the group size clusters (see Fig. S7 for species- and group-level results). Overall, there is a significant linear regression (β=0.718, r^2^=0.515, t_18_=4.37, p=0.0004). However, the data clearly form two distinct grades: the slope (and the goodness-of-fit) is lower in genera with large social groups (β=0.367, r^2^=0.135) than in those with smaller social groups (β=0.761, r^2^=0.580), suggesting that the upper grade relationship may be asymptotic. An asymptotic relationship would suggest a maximum *n*-clique size of ~9.0 at a degree size of ~4.0, with further increases in degree not yielding substantially larger *n*-cliques.

**Fig. 2.**
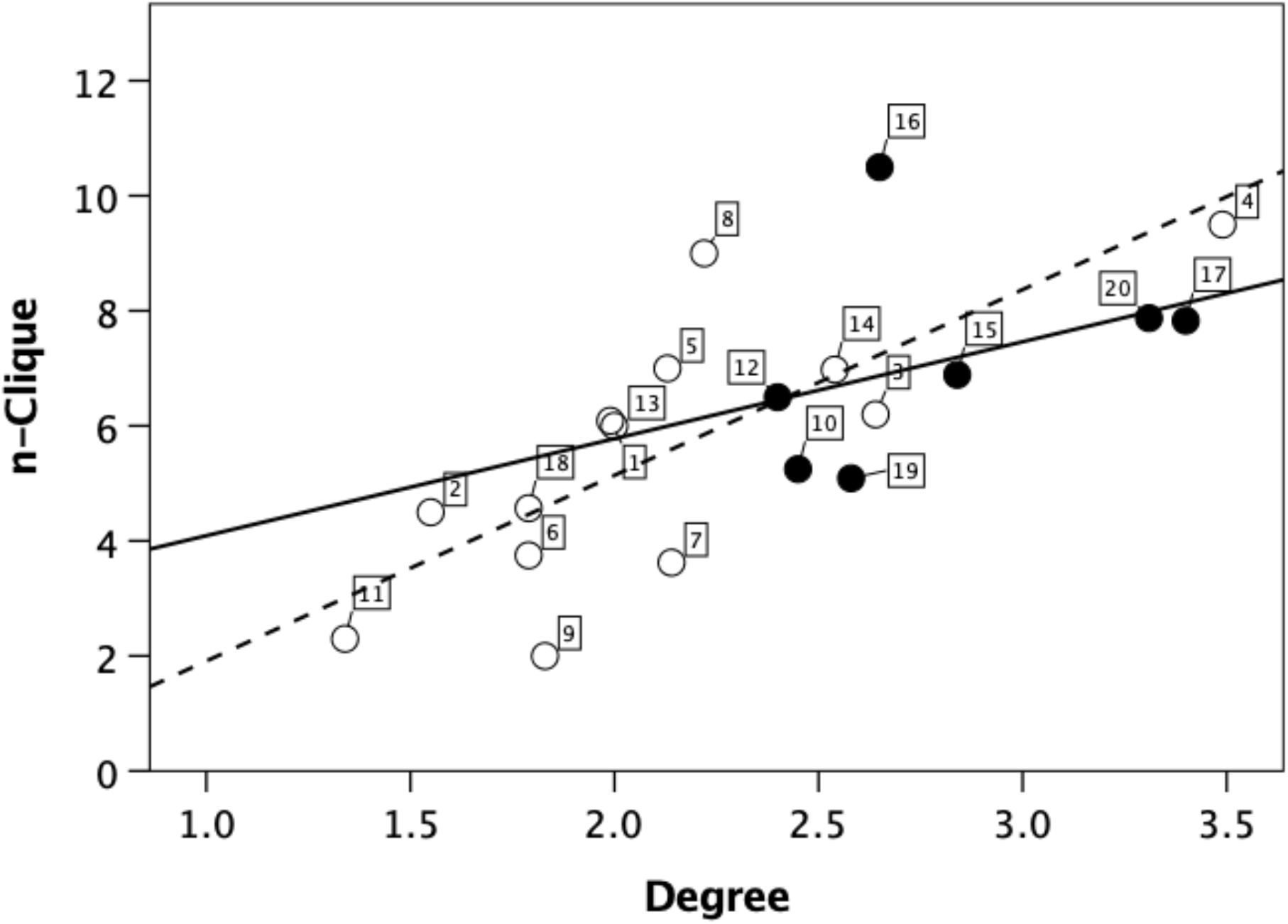
Mean n-clique plotted against mean degree for individual genera, differentiating genera with mean group size <25 (unfilled symbols, dashed regression) versus >25 (filled symbols, solid regression). Numbers attached to datapoints refer to taxa as listed in online *SI Dataset*.

Fig. 3 plots the mean number of adult females and mean total group size, respectively, against mean *n*-clique for each genus, differentiated by group size cluster. (Species and group-level plots are given in Figs. S8-S9, respectively.) It is obvious that the data form two parallel grades that differ in the number of reproductive females (12.4±3.1 versus 5.7±2.1: t_18_=8.43, p<0.0001) (Fig. 4). The mean ratio between the two grades is 2.4 for the number of females and 2.1 for total group size. The grades do not differ significantly in *n*-clique size (means of 6.81±1.9 versus 5.37±2.2: t_18_=1.21, p=0.242), but they do differ significantly in degree (upper grade species groom with more individuals: means of 2.68±0.6 versus 2.15±0.6; t_18_=2.16, p=0.044) (Fig. 4; see also Fig. S10).

**Fig. 3.**
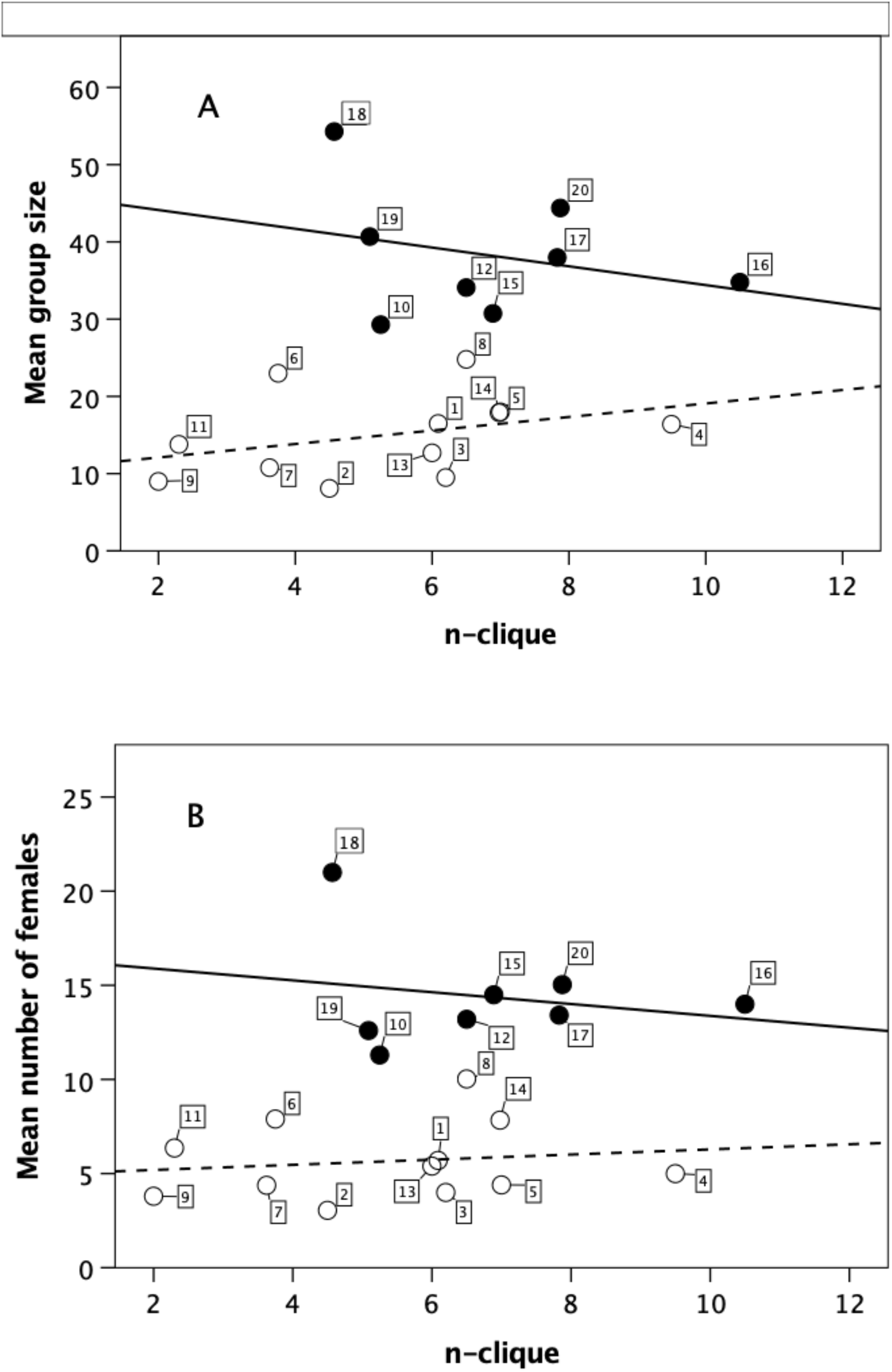
(a) Mean number of adult females per group and (b) mean group size for individual genera plotted against mean n-clique size, differentiating genera with mean group size <25 (unfilled symbols, dashed regression) versus >25 (filled symbols, solid regression). The social unit for *Theropithecus* is identified with the harem reproductive unit. Numbers attached to datapoints refer to taxa as listed in online *SI Dataset*.

**Fig. 4.**
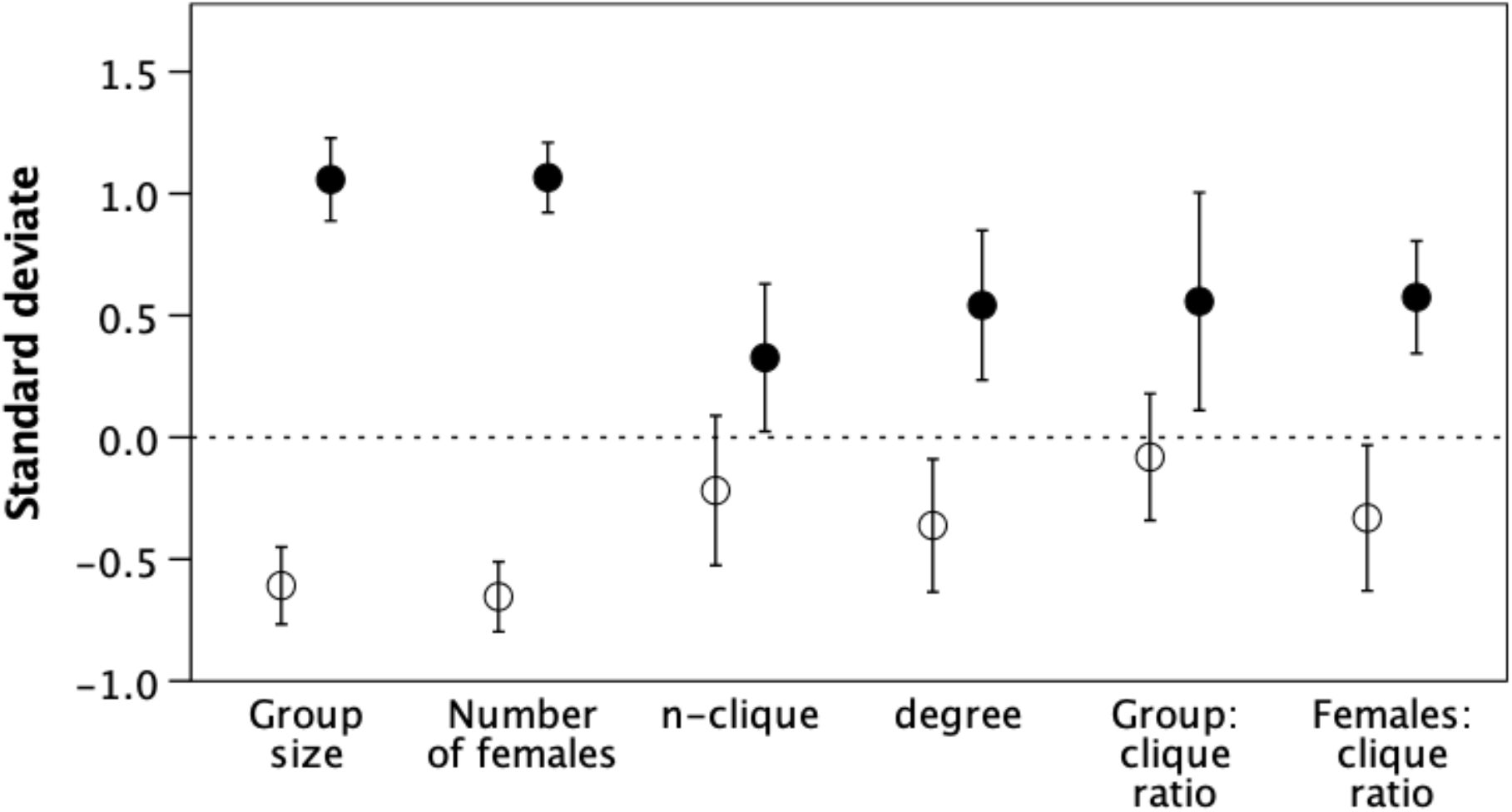
Mean (±1se) standard deviates for demographic variables, differentiating genera with mean group size <25 (unfilled symbols) versus >25 (filled symbols). The social unit for *Theropithecus* is identified with the harem reproductive unit.

The female:*n-*clique ratios (Fig. 4) are of particular significance. A ratio of ~1.0 would indicate that the number of females in the group is linearly determined by the size of the grooming chain: all females are included within a single *n*-clique; a ratio >1.0 would indicate that a chain typically includes only a proportion of the females, implying that there are fracture lines in the group’s grooming network that are fission risk points. In the lower grade, the female:*n*-clique ratios are not significantly different from unity (mean 1.21±0.69: t[ratio=1.0] = 1.27, df=11, p=0.247) indicating that most of the group’s females are included within a single grooming chain, whereas in upper grade genera the ratio approximates 2 (mean 1.78±0.48; t[ratio=1.0] = 4.92, df=8, p=0.001) indicating that the females are typically divided between two or more such chains. This implies that, in the upper grade genera, there are natural fracture points where the group is partitioned into two or more disconnected subnetworks, increasing the risk that groups will break up during foraging.

These data indicate that, although degree increases linearly with group size, extended grooming chains (*n*-cliques) do not correlate with the number of individuals who need to be bonded together in larger groups. More importantly, while small groups typically consist of a single *n*-clique, large groups consist of two (and hence are more at risk of fragmentation). Hypothesis H1 is supported only for small groups; this solution does not work for larger groups.

### Bonding (hypothesis H2)

Fig. 5 plots time devoted to grooming against group size and number of females. Time spent grooming increases exponentially with both group size (Fig. 5a: quadratic fit, r^2^=0.790, F_2,12_=22.5, p<0.0001) and the number of females (Fig. 5b: linear fit, r^2^=0.757, F_2,12_=18.7, p=0.0002). The upturn on the left hand side in Fig. 5a is due entirely to *Callithrix*. The callitrichids are unique among the primates in having a pairbonded mating system set within a larger group of adult and subadult helpers-at-the-nest. The higher rate of grooming may thus reflect the fact that bonding is especially intense in monogamous social systems.

**Fig. 5.**
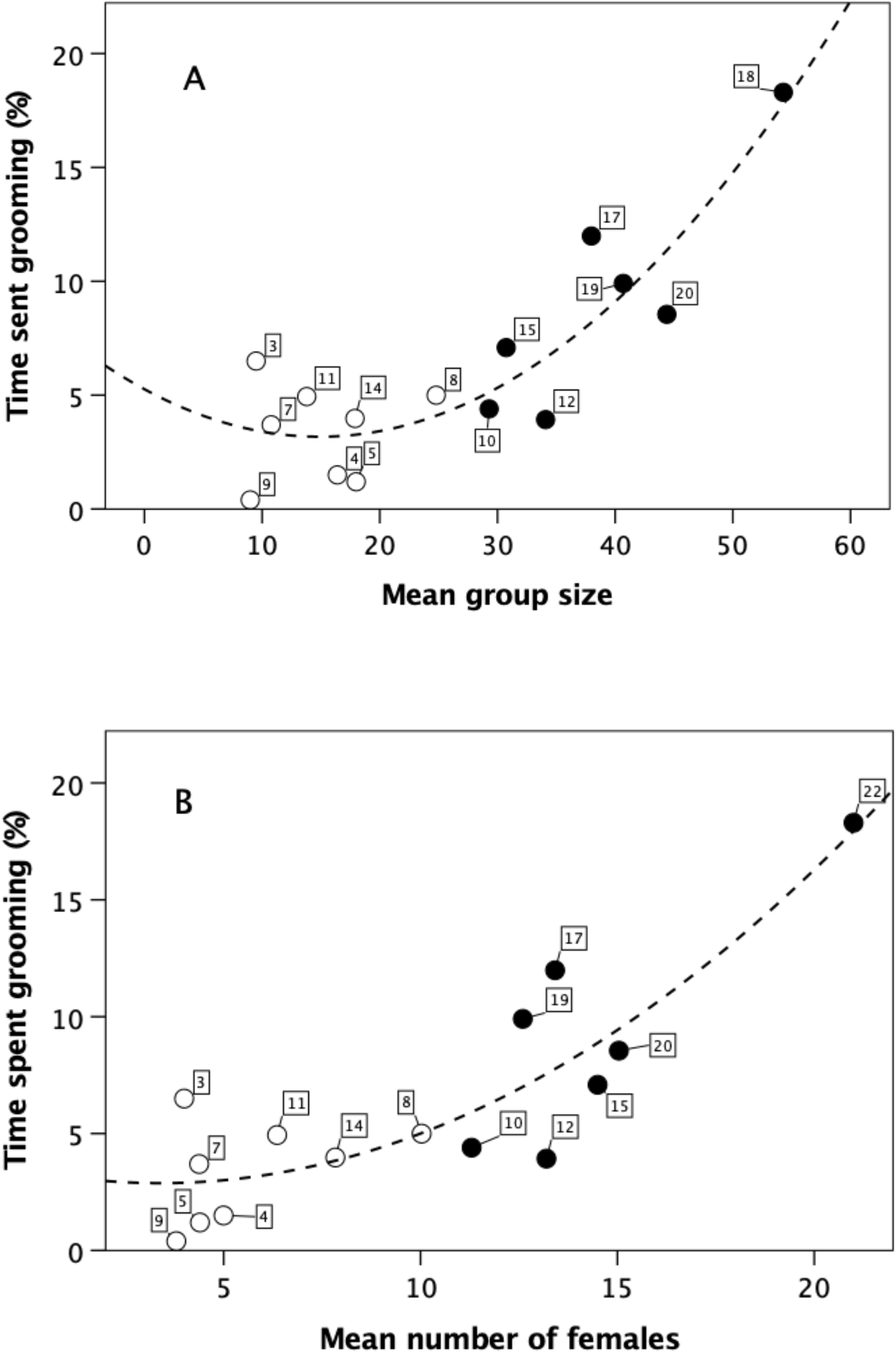
Time devoted to grooming for individual genera plotted against (a) mean group size and (b) mean number of adult females in the group, differentiating genera with mean group size <25 (unfilled symbols) versus >25 (filled symbols). The regression line has an exponential form. Numbers attached to datapoints refer to taxa as listed in online *SI Dataset*.

Animals’ time budgets are not infinitely flexible, and in many cases they have little spare time that can be devoted to grooming if the demand increases [45]. One way of solving this problem, however, might be to switch to a more frugivorous diet since fruits require less processing time than leaves and insects, and obviate the heavy investment in rest-based fermentation that leaves require [63]. Fig. 6 plots mean time that individual species devote to social grooming against the percentage of leaf in the diet. Although the available sample is small, it is clear that the pattern is very different for lower and upper grade taxa. For lower grade taxa, grooming time is not influenced by diet (linear regression: r^2^=0.053, p=0.620; cubic regression: r^2^=0.184, p=0.693), but for upper grade taxa there is a significant negative relationship (linear regression: r^2^=0.659, p=0.027; cubic regression: r^2^=0.864, p=0.003) that asymptotes at the same level as the lower grade taxa. A switch to an increasingly fruit-based diet frees off additional time from the foraging and resting time budgets to allow more time to be devoted to grooming.

**Fig. 6.**
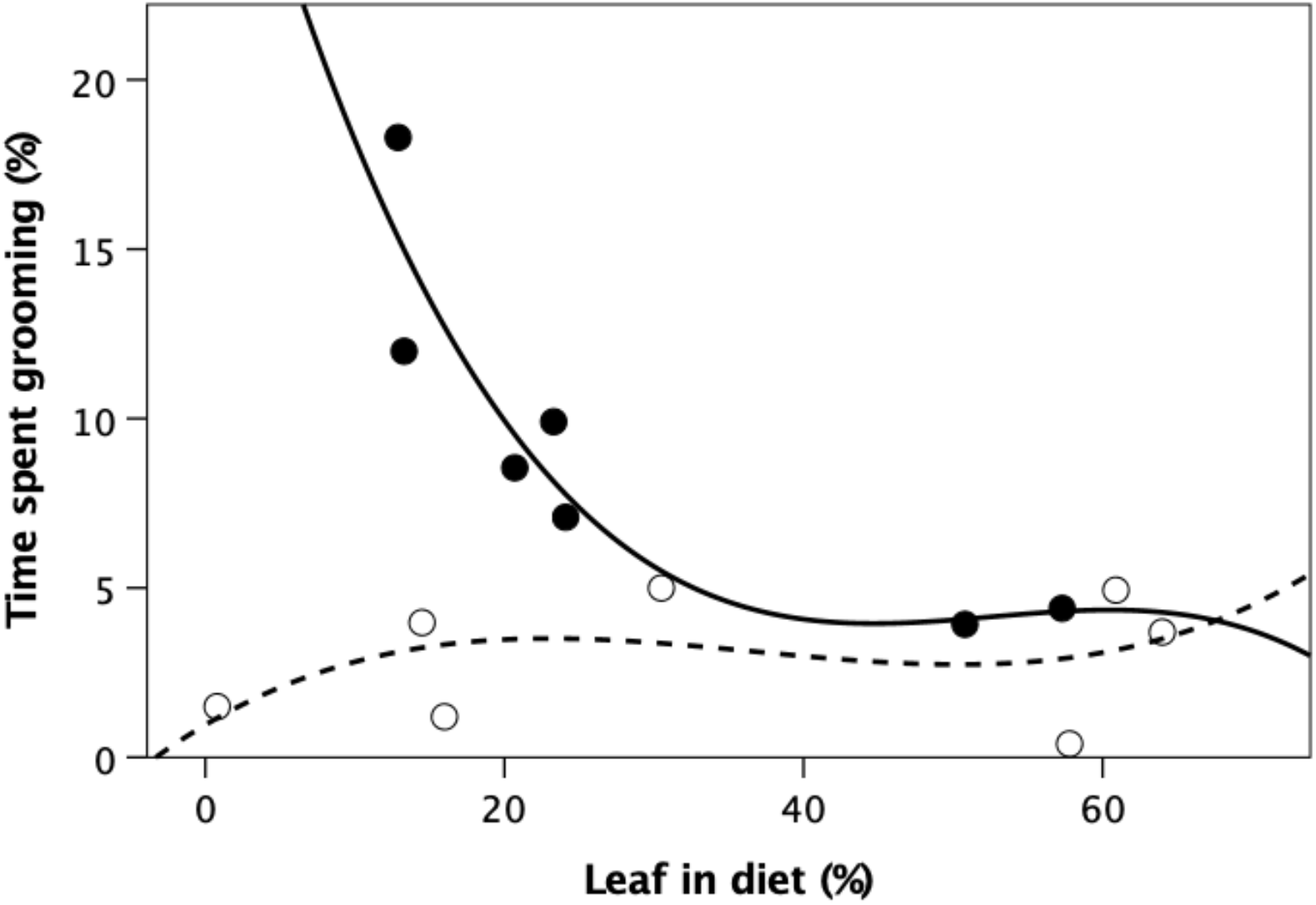
Mean time spent in social grooming (% of 12-hour day) as a function of the percentage of the diet composed of leafage for upper grade (filled symbols) and lower grade (unfilled symbols) taxa.

To test the directional hypothesis that upper grade taxa devote more time to social grooming, I compare values for total grooming time as well as grooming time per group member, per adult female and per degree for the two grades (Fig. 7a, Fig. S9). So they can be plotted on the same graph, I converted all scores to standard deviates from the respective overall mean for each index. The difference between the two grades in Fig. 7a is significant for total grooming (t_13_=3.00, p=0.005) and grooming time per degree (t_13_=2.16, p=0.025), but not for grooming time per group member (t_13_=-0.29, p=0.899) or per adult female (t_13_=0.14, p=0.495) (directional 1-tailed tests in each case). This suggests that, overall, upper grade genera increase time devoted to social grooming in proportion to group size, but that they invest the additional time disproportionally in their core grooming partners rather than spreading it equally around other members of the group. This suggests that, as group size increases, animals do seek to strengthen their core alliances, confirming hypothesis H2.

**Fig. 7.**
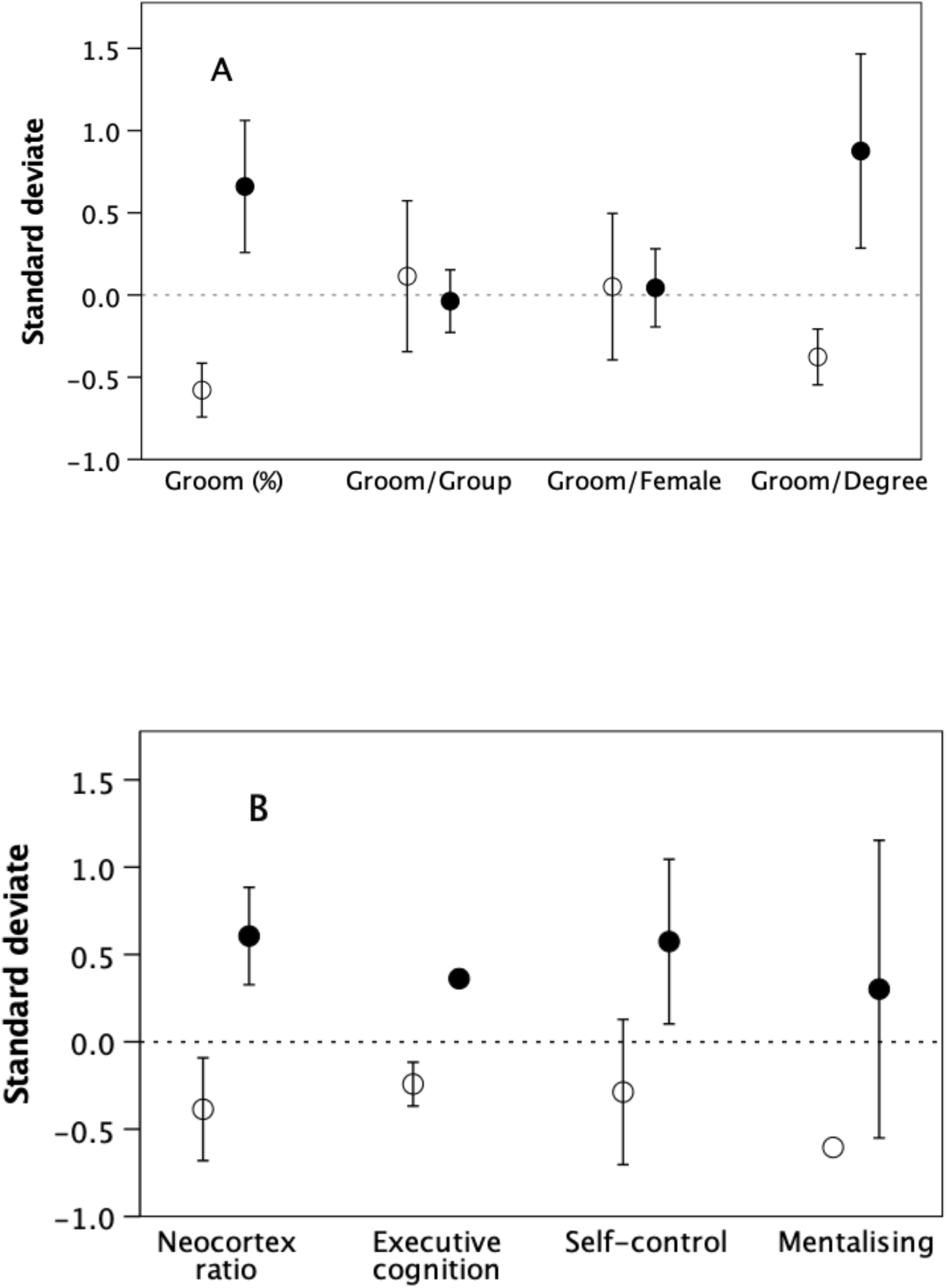
Mean (±1se) standard deviates for (a) social grooming indices and (b) four cognitive variables, differentiating genera with mean group size <25 (unfilled symbols) versus >25 (filled symbols). In (a), group size for *Theropithecus* is identified with the clan. In (b), neocortex ratio data are from [4]; executive cognition is the arcsin transform of the proportion of correct trials on 8 standard executive function tasks (from [52]); self-control is performance on an A-not-B task (from [53]); mentalising data are from [55].

### Cognition (hypothesis H3)

To test the directional hypothesis that upper grade genera have better social cognitive abilities, I compare performance on the four indices of cognition (neocortex ratio, executive cognition score, self-control and mentalising), with scores again transformed to standard deviates of the respective means (Fig. 7b). For all four indices, upper grade genera perform at higher levels than lower grade genera, though, due to small sample sizes, the differences are significant only for neocortex ratio and executive cognition (neocortex: t_16_=2.29, p=0.018; executive cognition: t_8_=3.74, p=0.003; self-control: t_7_=1.26, p=0.124; mentalising: t1=0.61, p=0.325) (directional 1-tailed tests in each case). Combining these tests using Fisher’s meta-analysis for small samples [65] yields χ^2^=26.09 (df=8, p=0.001), indicating that, overall, there is a consistent underlying trend for the upper grade genera to perform better than lower grade genera on all four indices. Hypothesis H3 is supported.

## Discussion

The results suggest that, in terms of network structure, primates divide into two distinct social grades: (1) genera that have small groups with fewer females, with grooming chains (*n*-cliques) that typically include all the adult females in the group and (2) genera that have large groups with more adult females and significant substructuring (more than one *n*-clique per group and a high female: *n-*clique ratio). Female:*n*-clique ratios that approximate unity (as in the lower grade) suggest that *n-*clique size imposes a limit on the number of females that can live together. This will inevitably limit group size [2]. The scaling ratio of ~2 for upper grade genera suggests that larger groups arise by postponing fission that, in the lower grade genera, would have occurred (and under conditions of low predation risk actually does occur even in upper grade genera [3,5]) at the size of the *n*-clique. The genera in the upper grade are all Old World monkeys and apes, but not all Old World monkeys are included in this grade. In fact, the taxonomic contrast between the two grades parallels that between weakly and strongly bonded monkey genera noted by [2,41]. As with the grades in the social brain relationship [41], these grades seem to reflect mosaic adaptation to contrasting ecological and demographic conditions, not phyletic grades.

In both grades, the size of the female cohort and total group size are an asymptotic function of extended grooming subnetwork (*n*-clique) size, suggesting that there are structural limits to the size of group that can be maintained as a coherent entity through personalised relationships. Groups can become more substructured, but the resulting subnetworks cannot contain more than the asymptotic number of females. This suggests that only certain group sizes are possible [3,41]. Fig. 3b suggests that the limits on the number of females in the two grades are ~6 and ~14, respectively. This is in close agreement with the finding that, in primates, fertility is optimised at female cohort sizes of 6.9 and 13.6 in the two grades that correspond to those in Fig. 3b [2]. These equate to limits on group size of ~18 and ~37 (Fig. 3a), close to the previously identified attractors for group size at ~16 and ~31 [3].

The central issue, then, is how upper grade genera prevent their groups fracturing at the size at which this happens in the lower grade genera. The fact that *n*-cliques do not include all adults (or even all females) rules out the structural hypothesis (H1) that grooming chains bind all individuals into a single group. Since both hypotheses H2 and H3 are supported, this suggests that this is achieved by behavioural (bonding) and cognitive mechanisms, probably being introduced stepwise during the course of evolution. Genera of the upper grade disproportionately increase the amount of time devoted to grooming their core social partners as group size increases (Figs. 5, 7a), providing they can effect change in diet to free of sufficient time to do so (Fig. 6). This provides an effective buffer against the stresses created by the proximity of many other individuals [2], not least by the simple device of causing others to maintain their distance rather than risk being attacked by several individuals. Additional support for this is given by the fact that, in primates, rates of conflict within female dyads are negatively correlated with species group size when controlling for neocortex ratio [65]. Grooming is the basis of coalition-formation in primates [24,49,66], and it is conspicuous that formal coalitions are universal in all the upper grade genera but rare (if not completely absent) among lower grade genera. The capacity to form coalitions is what makes it possible for the upper grade genera to defer the stress-induced infertility effects so as to live in larger groups [2].

However, while coalitions solve the stress problem, they don’t solve the associated coordination problem. Something else is needed to counteract the natural fragmentation process so that groups remain cohesive over time despite pressures favouring dispersal. That something seems to involve the capacity to make rapid inferences about high level rules (one-trial learning), understanding relationships beyond the confines of one’s own immediate grooming circle and what amount to the skills of diplomacy (knowing when to escalate a conflict and when not). These involve the ability to make rapid judgments about the meaning of signals and the nature of third party relationships so as to avoid escalating conflicts unnecessarily.

Understanding third party behaviour has been reported for many of the upper grade genera (*Pan* [67–69]; *Papio* [70–71]; *Macaca* [72–74]), but convincing evidence for such behaviour has not so far been reported for any of the lower grade genera. Similarly, many upper grade genera are able to evaluate the status of another individual simultaneously on two or more separate dimensions (e.g. kinship versus rank: *Papio* [75]; *Macaca* [76]), whom another individual has alliances with (*Pan* [67]; *Papio* [70,77]; *Macaca* [78–79]) and, on the basis of observed reputation, how trustworthy they might be (*Pan* [68]). Again, such competences have not, so far, been reported for any of the lower grade genera. Reconciliation (repairing relationships destabilised by conflict), a behaviour that depends on the ability to recognise that a relationship has been weakened (thus implying some minimal capacity to mentalise), has been widely reported from upper grade genera but rarely (and usually with mixed results and only in the form of physical proximity without involving conciliatory signals or active grooming) in lower grade genera [80].

The finding that cognitive abilities and relative neocortex size seem to play a central role in managing relationships is reinforced by evidence from neuroimaging studies. In both monkeys (*Macaca* [81–82]; *Papio* [83]) and humans [84–88], the size of personal social networks correlates with the volume of the brain’s default mode network and its extensions down into the limbic system and the cerebellum. In primates, this very large connectome, with its massive white matter connections, takes up a substantial proportion of the non-visual cortex, contributing significantly to overall brain size. Moreover, it exhibits significant enlargement in anthropoid primates compared to prosimians [89].

In sum, it seems that the ecological need to live in large, stable social groups has necessitated finding ways to mitigate the escalating effects of stress so as to reduce the risk of groups fragmenting. This seems to have required a combination of investing more heavily both in additional grooming time for enhanced bonding of core alliances to mitigate the direct costs of stress and in social cognitive skills that are dependent on specialised neural circuits in the brain for managing relationships beyond this inner circle. This distinction between close-bonded and peripheral relationships within a network is reminiscent of Grannovetter’s [90] concept of weak and strong ties in human social networks. These findings also provide a *prima facie* case for the claim that primate species that live in large social groups exploit more sophisticated forms of cognition, and hence why they should have evolved much larger brains than other orders. Nonetheless, although primates are, as a group, the most social of all the mammals, the same issues apply broadly to all social mammals. Understanding how other taxa that live in bonded social groups (equids, tylopods, elephants, delphinids, sciurids) solve these same coordination problems would add measurably to our understanding of the processes of social evolution.

## Supporting information

Structural and Cognitive Solutions to Prevent Group Fragmentation SI

## Notes

### Competing Interest Statement

The authors have declared no competing interest.

