## Supplementary material for "Structural and Cognitive Solutions to Prevent Group Fragmentation in Group-Living Species": Structural and Cognitive Solutions to Prevent Group Fragmentation SI

**R.I.M. Dunbar**

### *Supplementary Information*

#### Supplementary Methods

*Defining social bonds and social groups*

Many studies (Silk et al. 2013; Pasquaretta et al. 2014; Duboscq et al. 2016) have used spatial proximity as a metric for defining affiliative networks. When the focus of a study is on the behavioural *consequences* of relationships (e.g. information or disease transmission), networks defined in this way are appropriate. Animals can, quite obviously, infect each other or learn from each other merely by being in physical proximity; they do not need to groom to do so. However, this is *not* the case when the focus is on how relationships are built and maintained (the emotional content of relationships associated with bonding): proximity (i.e. tolerance) is a *consequence* of having a relationship, not the way the relationship is created. More importantly, networks based on different criteria (e.g. proximity and grooming) do not always correlate especially well because they are measuring very different social phenomena (Silk et al. 2007; Castles et al. 2014; Smith-Aguilar et al. 2019). Since, for the reasons indicated in the main text, grooming is central to the creation and maintenance of bonded relationships in primates and our concern is specifically with the effort individuals invest in creating relationships, I consider only networks based on social grooming. For present purposes, I am interested only in relationships that involve adults and post-puberty subadults. Immatures typically make up ~50% of a primate group (Dunbar et al. 2018). However, socially, most immatures are simply appendages of their mothers and rarely groom much with anyone else (orphans, for example, are invariably social isolates). Grooming with or between pre-puberty immatures was not, therefore, counted.

For a study to be included in the sample, I required either a matrix of grooming frequencies or a weighted sociogram specifying at least the relative frequencies of grooming between all adult individuals. Because animals occasionally groom with individuals outside their immediate grooming cliques, networks quickly become saturated (everyone grooms with everyone else) (James et al. 2009). Relationship quality is a function of the time invested in it in both monkeys and humans (Dunbar 1980; Roberts & Dunbar 2011; Sutcliffe et al. 2012), not a consequence of the fact of interacting just once.

For most species of primates, the identity of the group is uncontroversial: the animals forage and travel as a single, defined set of animals whose composition remains stable (subject to births and deaths) over considerable periods of time. However, in species with multilevel social systems, foraging and social groups have different sizes, the former being a subcomponent of the latter. These come in two forms: atomistic systems (represented by chimpanzees) and modular systems (represented by gelada). Chimpanzees forage in small parties (typically 3-5 individuals) of flexible membership that form a single, demographically stable community that occupies a defined territory (Dunbar et al. 2018). The gelada (*Theropithecus*), in contrast, live in small, stable reproductive harems (of 5-25 individuals) that forage alone or in herds that vary in size between 20-300 (2-30 harems); harems typically form herds only with units that belong to the same higher level groupings (teams, bands and communities: Kawai et al. 1983) whose membership is consistent through time. This structural diversity makes it especially important to determine the correct level for any given analysis given the functional context of interest. Because the clan is a stable association of 2-5 harems that is, functionally, equivalent to a *Papio* troop (MacCarron & Dunbar 2016), I use the clan for gelada.

*Defining weak vs strong ties*

To avoid saturated networks where everyone grooms with everyone else, I follow standard practice and use a cut off value defined by the mean frequency of grooming per available dyad assuming animals distributed their grooming effort at random to all possible alters:

$$\text{Mean tie strength} = \Sigma\text{Gm}/0.5(\text{M}^2\text{-M})$$

where $\Sigma$Gm is the total number of grooming bouts observed and *M* is the total number of mature (i.e. adult or subadult) animals in the group. I use undirected matrices (i.e. the data do not distinguish direction of grooming) since we are not specifically interested in who is *responsible* for maintain the relationship, only whether or not a relationship exists.

In analyses of network structure, it is conventional to distinguish between casual (weak) and substantive (strong) ties (Granovetter 1973). Typically, in most species, individuals have a few strong ties and many casual ties (Sueur et al. 2011; Sueur & Maire 2014; Sutcliffe et al. 2012). Casual ties commonly have high turnover, whereas more substantive relationships typically have considerable temporal stability (Duboscq et al. 2016; Roy et al. 2022). Since we are interested in longlasting relationships, we seek a rule that allows us to differentiate between weak and strong ties.

Most distributions of this type follow a power law (Fig. S1). Under a conventional 'broken stick' model, the criterion for distinguishing between weak and strong relationships is the point at which the graph changes pitch (indicated by the dashed vertical line). In each case, this point is very close to both the point at which the slope first changes direction (the inflexion point) and the mean value across all dyads (the 50$^{th}$ centile). For an asymptotic distribution, the inflexion point is given by the point on the X-axis that corresponds to the point that is 1/$e^{th}$ down from the asymptotic value on the Y-axis (indicated by the horizontal line).

The three samples in Fig. S1 all use different metrics based on grooming frequencies, but all seem to be in close agreement: there is a clear distinction between weak and strong ties, and the cut-off point lies just above the mean value (defined by the 50$^{th}$ centile). Since all these indices are in close agreement, and I use grooming frequencies for all the datasets, I use the mean frequency with which dyads would groom each other if all the observed grooming sampled from a group is distributed equally between all possible dyads (the mean rate per possible dyad). In those cases where the data are in the form of a sociogram that distinguishes strength of the tie in to a number of categories that differ in strength, I placed the cut-off between the two lowest bands (i.e. the lowest band was excluded) for cases that specified only 4 gradations or after the lowest two bands if there were more gradations.

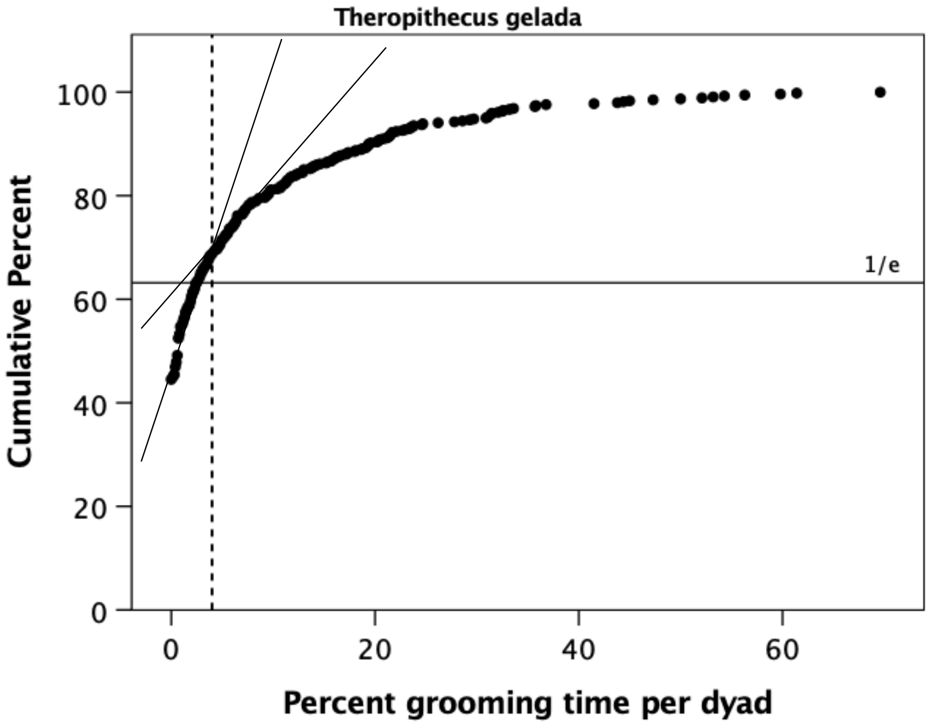

Theropithecus gelada
Cumulative Percent
Percent grooming time per dyad
1/e
0
20
40
60
80
100
0
20
40
60


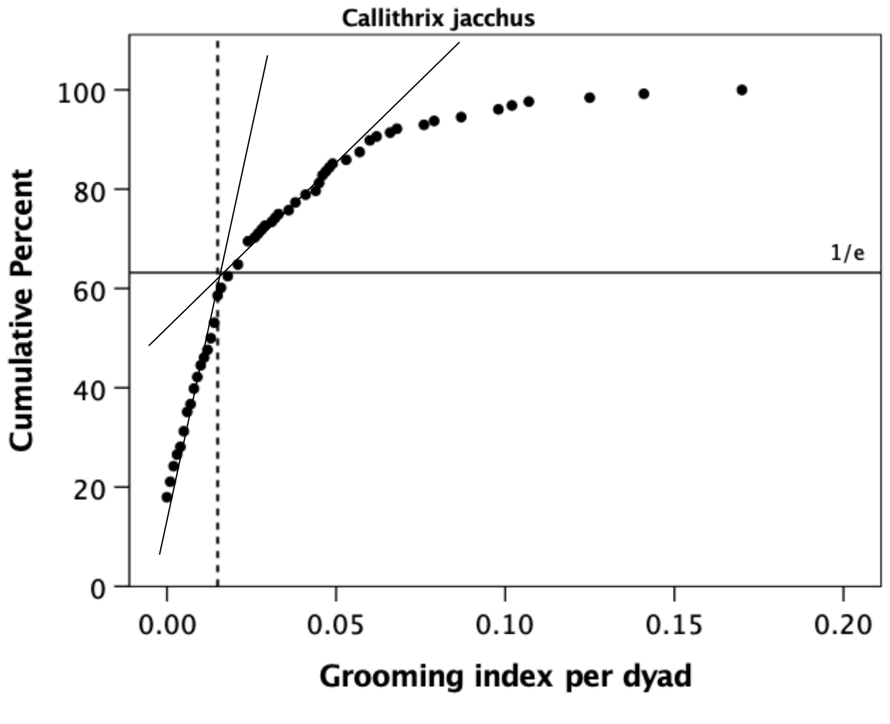

Callithrix jacchus
Cumulative Percent
Grooming index per dyad
1/e
0
20
40
60
80
100
0.00
0.05
0.10
0.15
0.20

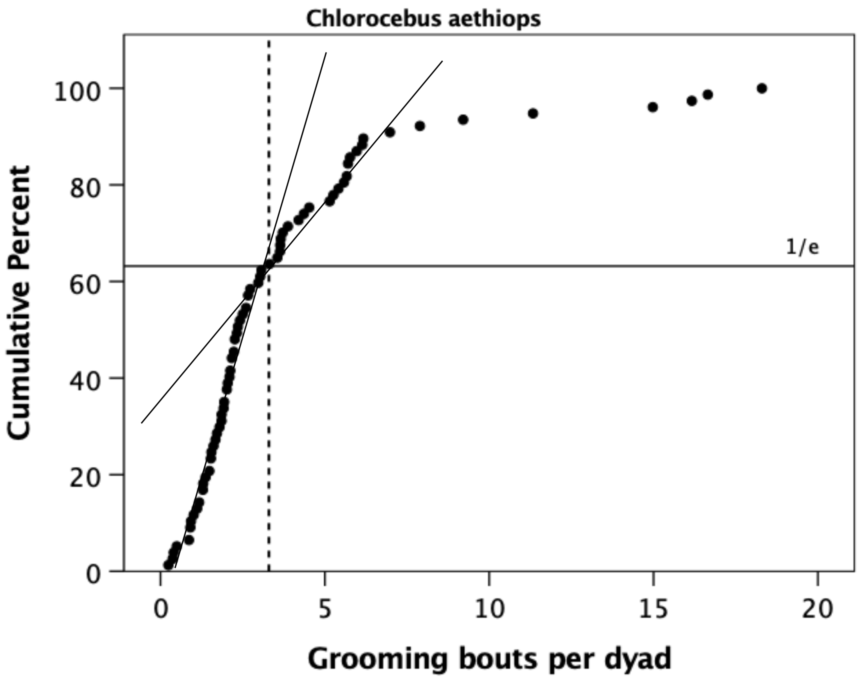


***Figure S1***

*Cumulative distribution of grooming effort in individual dyads for three different species, based on different metrics. In each case, the dashed vertical line demarcates the point at which the graph changes pitch defining a difference between weak and strong relationships under a conventional 'broken stick' model (indicated by the pairs of thin lines); the horizontal line identifies the point that is 1/e$^{th}$ down from the asymptote (defining the point at which the slope changes). In each case, this is close to the 50$^{th}$ centile (the mean value). Sources:* T. gelada*, R. Dunbar (unpublished data);* C. jacchus*, Lazaro-Perea (2000);* C. aethiops*, Seyfarth (1980).*

*Optimal number of clusters*

For this analysis, I used the species-level data since it provides a larger sample. However, an analysis of the genus-level data yields identical results despite the smaller sample. I give three different ways of identifying the optimal partition.

Figure S2 uses the classic 'broken stick' method used extensively in behavioural ecology to identify a criterion for bout length. It seeks the point where there is a change in slope in the cumulative distribution, and takes the value(s) on the X-axis as the break-point(s) to differentiate subsets, or clusters, within the data. The point where they cross demarcates the optimal cut-off, and it is at a group size of N=27.5.

A second method partitions the data stepwise across the range of the X-axis and determines the goodness of fit when separate regressions are placed through the lefthand and righthand data subsets. We seek the value of X where goodness-of-fit reaches an asymptote. Fig. S3 plots the goodness-of-fit against different cut-off values for mean species group size: fit is maximised at N≈25.

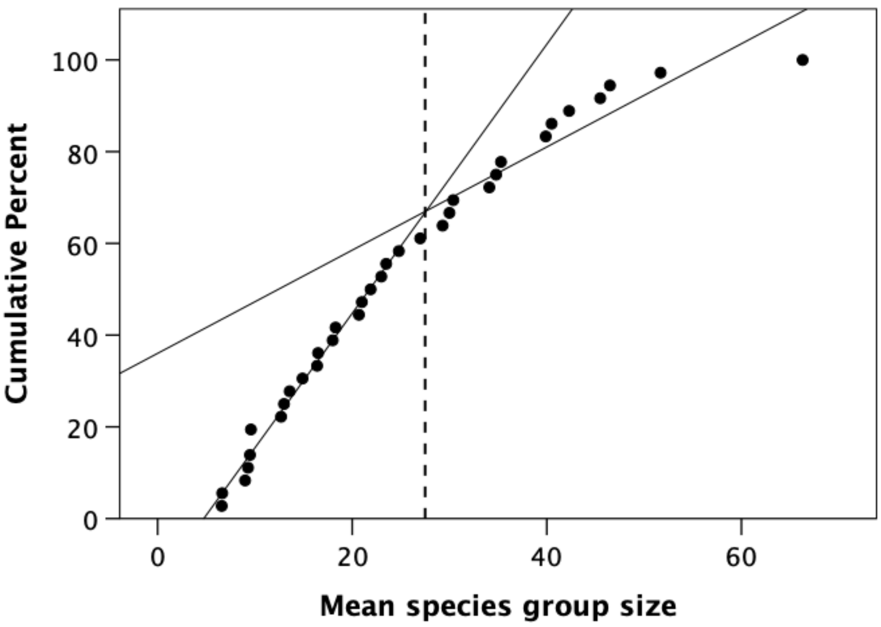


***Figure S2***

*Cumulative percentage distribution of mean social group size for individual species, with a 'broken stick' analysis (narrow solid regression slopes) to identify the optimal partition into two main clusters at a group size of N=26 (vertical dashed line).*

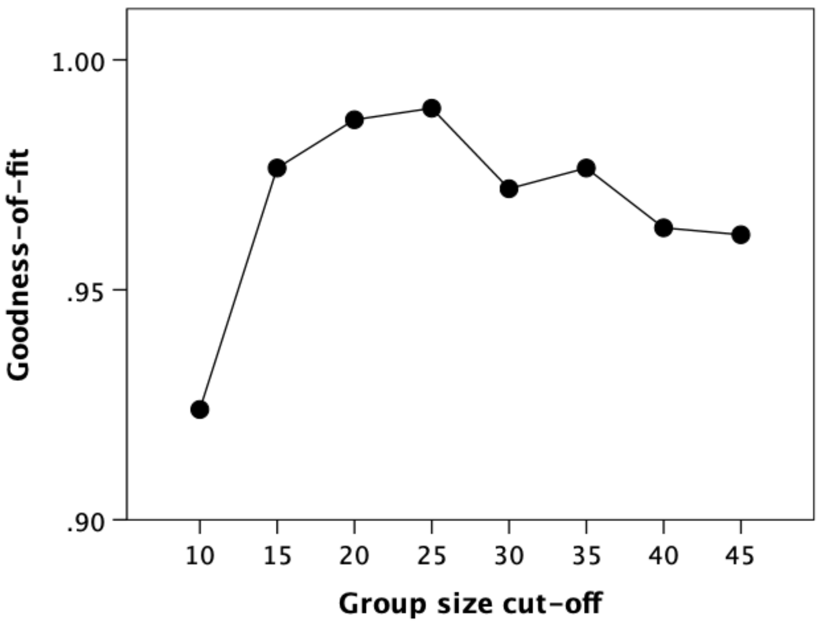


***Figure S3***

*Goodness-of-fit (indexed as $r^2$) for different cut-off points for the cumulative distribution of group sizes shown in Figure S2. For calculating regressions, the outlier point on the extreme right of Figure S2 was discounted.*

A third approach uses *k*-means cluster analysis to find the optimal way to partition the data in terms of the distribution of mean group size. We seek the minimum number of partitions that maximises goodness of fit, where the distribution across clusters is not too uneven and no cluster contains only a single value. I considered partition into *k*=2-7 clusters. Figure S4 plots the clusters identified for values of *k*=2-5. The superimposed plots of the partitions suggest that there is a primary division into two main clusters (demarcated in red), with values of *k*>2 simply subdividing these two partitions in different ways. A *k*=2 solution gives a significant partition ($F_{1,34}$=84.57, p<0.0001). The fit is slightly poorer (but still significant) at *k*=3 ($F_{2,33}$=71.53, p<0.0001), but improves thereafter. However, all partitions for *k*>2 include at least one cluster with a single member (usually considered undesirable). The *k*=2 solution partitions the data at a group size of N=28 into clusters with 28 and 14 species. The equivalent values for a genus-level analysis are two clusters of 12 and 8 genera ($F_{1,18}$=59.36, p<0.0001) and a partition at N=27, with all analyses for *k*>2 yielding at least one cluster with a single member.

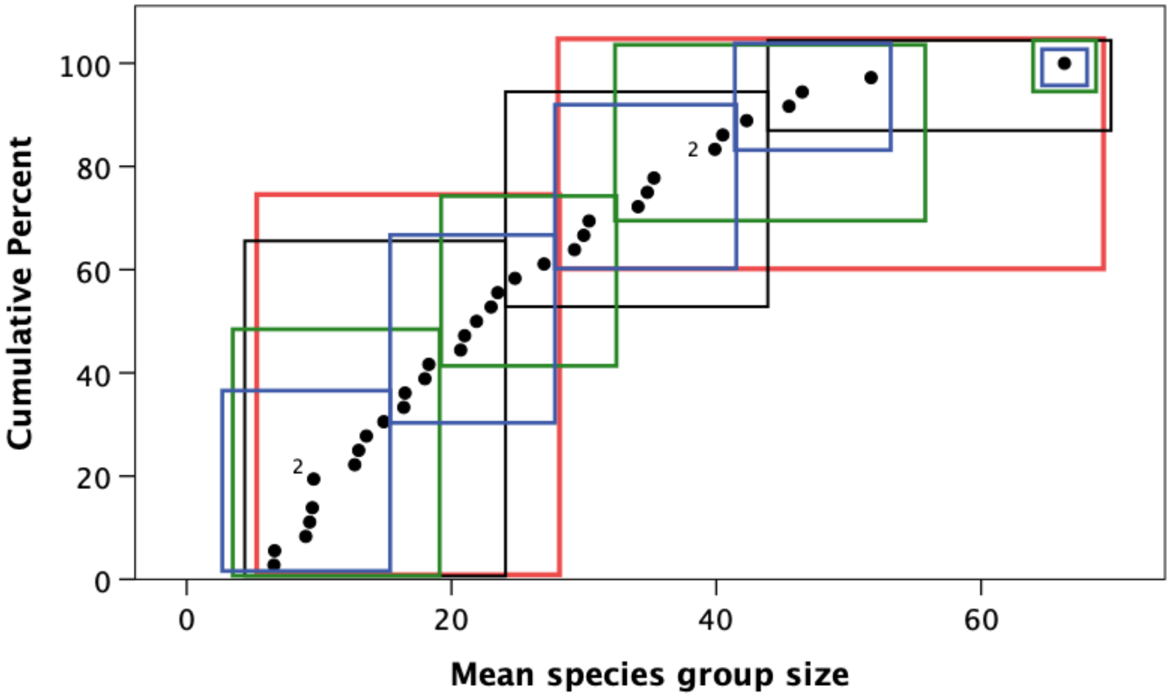


**Figure S4**

*Cumulative percentage distribution of mean social group size for social species, with optimal division into clusters by k-means cluster analysis for k=2 (red), k=3 (black), k=4 (green) and k=5 (blue). Superscript numbers indicate two superimposed datapoints. Partitions into k=3-5 clusters basically subdivide the k=2 division rather than creating novel ways of partitioning the data. The dashed purple box indicates the main overlap zone centred on N≈28.*

The three methods broadly agree that, for this sample of species, the group size data partition into two clusters, giving close agreement with Dunbar et al. (2018). The three methods give very similar estimates of where the optimal partition lies (range 25-28). Since the mean group size is 25.3 for the species-level data (with no species having a group size in the interval 24.8<N<27) and the data exhibit a clear bimodal distribution (Figure S5), I will take N=25 as the criterion for dividing species into those that live in small and large groups.

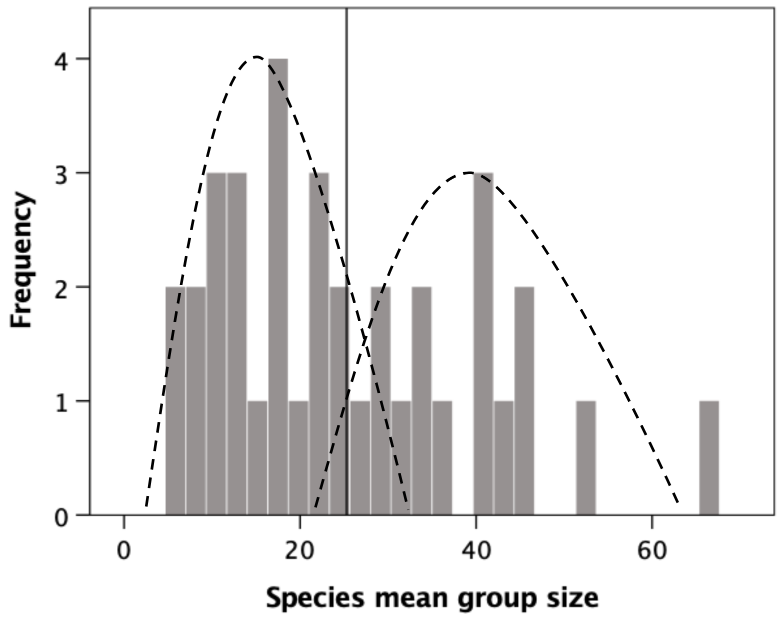


**Figure S5**

*Frequency distribution for individual species mean group size values for the sampled species. The vertical solid line demarcates the sample mean* (*25.3*)*; the dashed lines are the normal distributions for the two subsamples partitioned at the mean* (*i.e. for species with group sizes <25.3* [*mean size 15.2*] *and those with group sizes >25.3* [*mean size 39.6*]).

*Phylogenetic autocorrelation*

As a check that phylogenetic autocorrelation does not yield specious results due to inflated degrees of freedom, I ran the main analyses with phylogenetic correction using the consensus primate tree downloaded from https://10ktrees.nunn-lab.org/. The results are shown in Fig. S6 for the analyses shown in Figs. 2 and 3a. In neither case are the results different to the analyses without phylogenetic correction (regressions set through the origin: n-clique/degree, $r^2$=0.439, $t_{17}$=3.75, p=0.001; group/n-clique, $r^2$=0.443, $t_{17}$=3.78, p=0.001). Identifying grades is not straightforward with contrasts, in part because contrasts analysis overrides grade differences (unless they are strictly taxonomic, in which case they will appear as an obvious outlier) and in part because the plotted data are comparisons between nodes not individual taxa. Nonetheless, the wide tube-like distribution of the data in Fig. S6b compared to that in Fig. S6a (which is bivariate normal) is suggestive of the presence of grades.

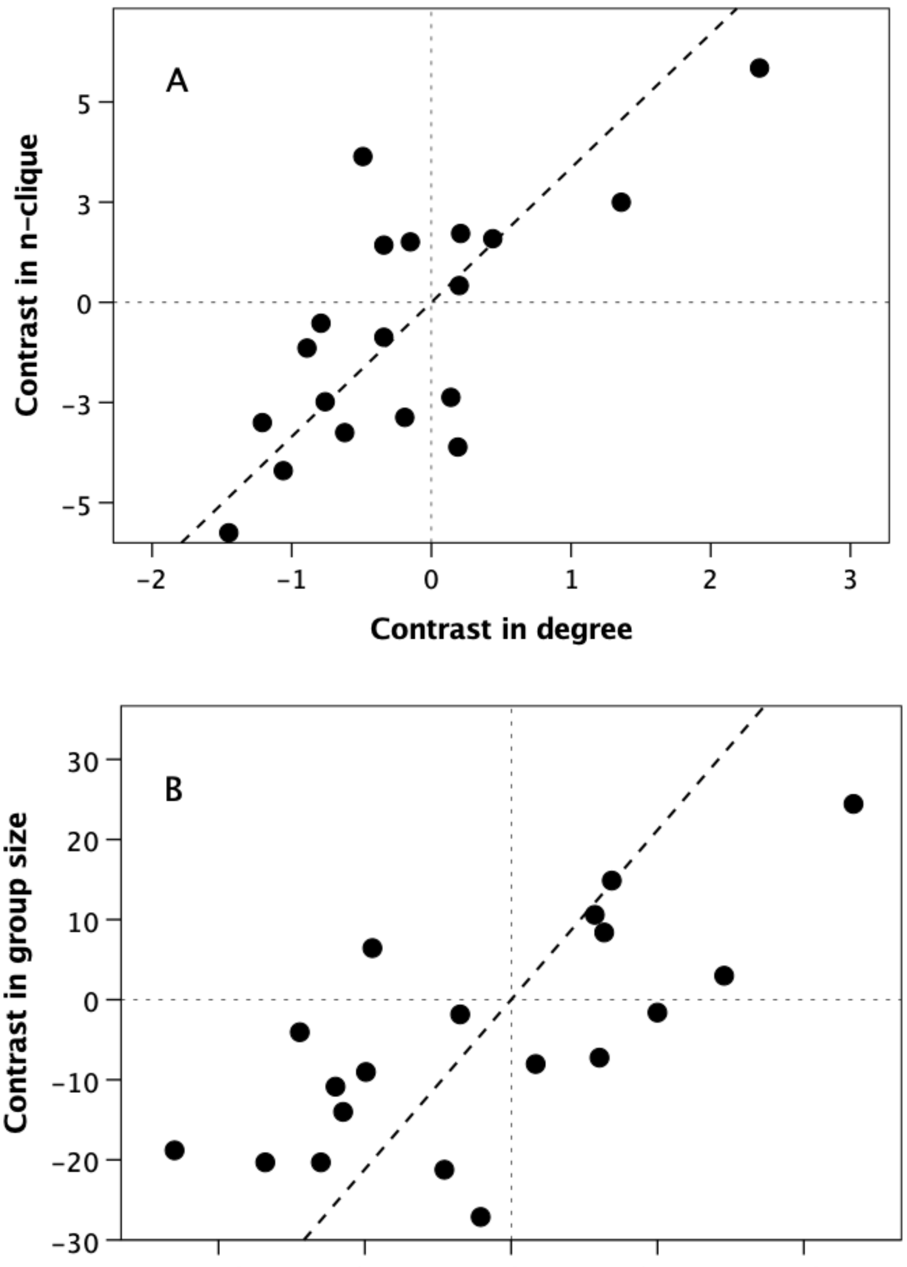


***Figure S6***

*Phylogenetically controlled plots of mean genus values for (a) n-clique (network) size plotted against degree (grooming clique size) (OLS regression set through the origin: $t_{18}$=3.75, p=0.001) and (b) group size plotted against n-clique size ($t_{18}$=3.78, p=0.001). Regression lines are set through the origin, as conventional.*

**Supplementary results**

Figures S7-S10 give the equivalent results to those in Figs. for the data at the species (plots [A] and group [plots [B] levels.

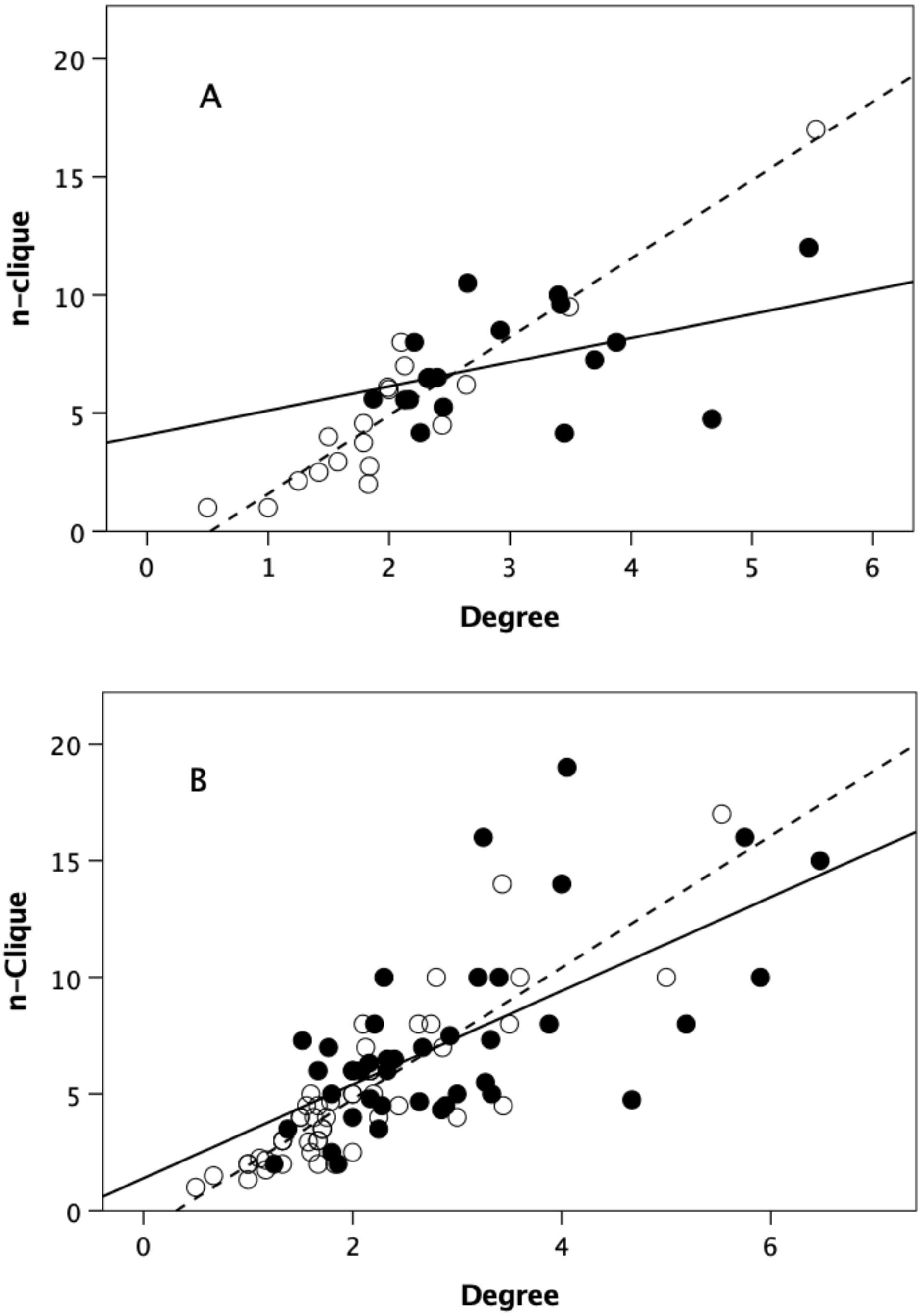


***Figure S7***

*Mean* n*-clique plotted against mean degree for (a) individual species and (b) individual groups in the sampled populations. Filled symbols and solid line: genera with group size >25; unfilled symbols and dashed line: genera with group size <25. Overall OLS regressions: (a) $\beta$=0.780, $r^2$=0.608, $t_{34}$=7.27, p<0.0001; (b) $\beta$=0.776, $r^2$=0.587, $t_{89}$=11.28, p<0.0001. The asymptotic character of the relationship is especially clear in Fig. S6(a).*

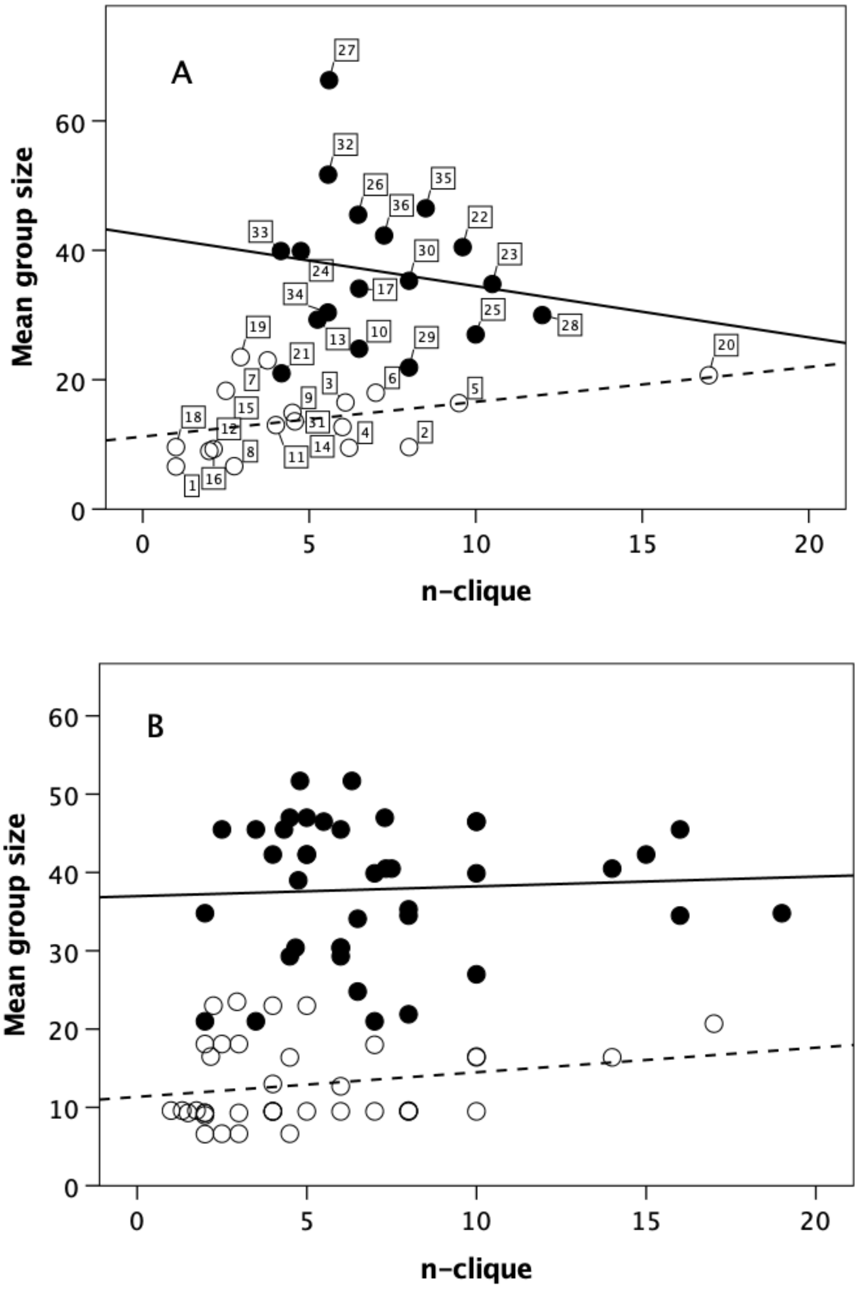


***Figure S8***

*Mean group size plotted against mean n-clique for (a) individual species and (b) individual groups in the sample in the sample populations. Filled symbols and solid line: genera with group size >25; unfilled symbols and dashed line: genera with group size <25. The two grades differ significantly: (a) means of 37.7±8.2 versus 13.9±5.3, respectively: $t_{34}$=7.71, p<0.0001 and (b) means of (37.9±8.7 versus 23.5±18.8: $t_{90}$=4.75, p<0.0001).*

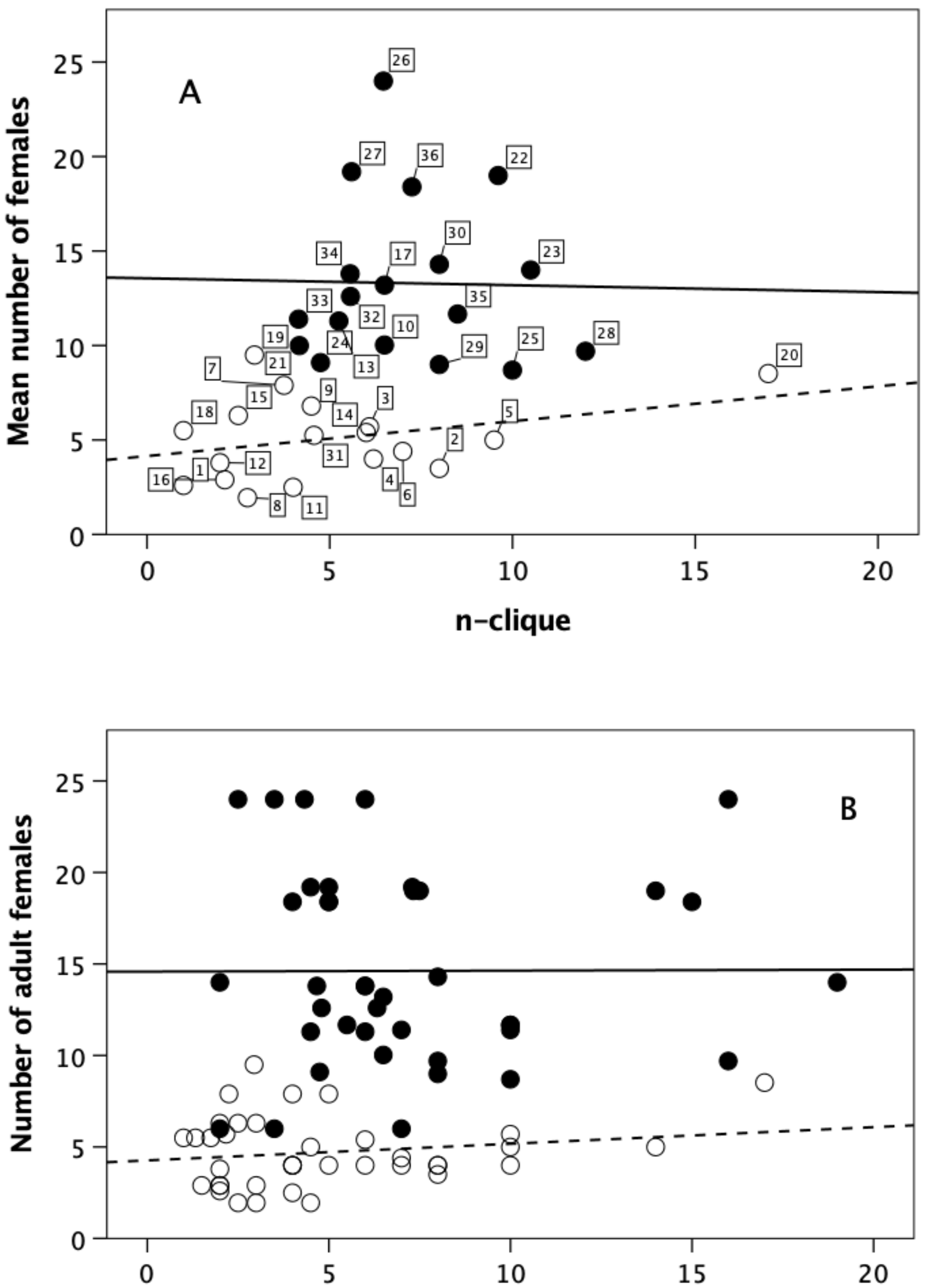


***Figure S9***

*Mean number of adult females per group plotted against mean n-clique for (a) individual species and (b) individual groups in the sample. Filled symbols and solid line: upper grade genera (as defined by genus-level analyses); unfilled symbols and dashed line: lower grade genera. The two grades differ significantly: (a) means of 14.5±5.2 versus 8.9±7.4, respectively:* $t_{34}=4.56$, $p<0.0001$ *and (b) means of (14.5±5.2 versus 8.9±18.8:* $t_{90}=4.75$, $p<0.0001$*).*

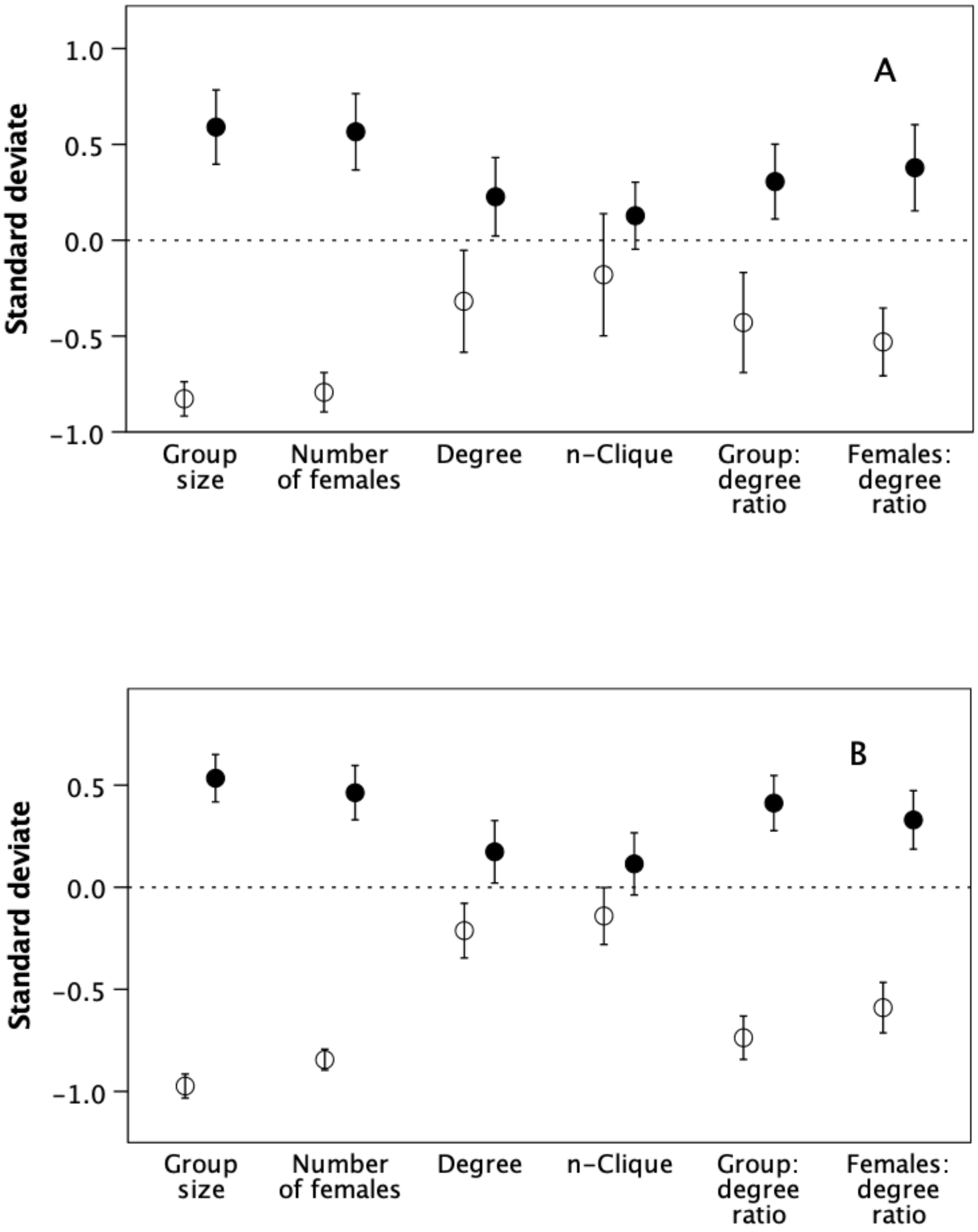


***Figure S10***

*Mean±1se for demographic variables for (a) species and (b) individual groups.*
*Filled symbols: upper grade genera (as defined by genus-level analyses); unfilled symbols: lower grade genera.*
